## Supplementary Information for "A higher order PUF complex is central to regulation of *C. elegans* germline stem cells"

**Supplementary Table 1.** Crystallographic data collection and refinement statistics.

| | **FBF-2 RBD/FBEa*** |
|---|---|
| Resolution range (Å) | 20.0-2.29 (2.33-2.29)[1] |
| Space group | $P6_1$ |
| Unit cell dimensions<br>a, b, c (Å)<br>α, β, γ (°) | <br>99.0, 99.0, 107.2<br>90, 90, 120 |
| Unique reflections[2] | 26610 (1305) |
| Multiplicity | 7.3 (7.4) |
| Completeness (%) | 99.9 (99.9) |
| Mean I/sigma(I) | 13.5 (2.0) |
| Wilson B-factor | 50.7 |
| R-meas | 0.15 (0.80) |
| R-pim | 0.06 (0.29) |
| **Refinement** | |
| Reflections used in refinement | 26063 |
| Reflections used for R-free | 1963 |
| R-work | 0.191 (0.257) |
| R-free | 0.232 (0.299) |
| Number of atoms | |
| protein | 3157 |
| RNA | 212 |
| solvent | 89 |
| RMSD bonds (Å) | 0.002 |
| RMSD angles (°) | 0.371 |
| Ramachandran favoured (%) | 98.2 |
| Ramachandran outliers (%) | 0 |
| Average B-factors ($Å^2$) | |
| protein | 65.2 |
| RNA | 79.1 |
| solvent | 58.3 |

[1]The highest-resolution shell is shown in parentheses.

[2]Statistics for the highest-resolution shell are shown in parentheses.

**Supplementary Table 2.** Statistical significance for GLD-1 quantitation comparisons

| Figure | Strains compared | Region (µm from distal end) | p-value | Sig[1] |
|---|---|---|---|---|
| | **Endogenous** | | | |
| 3D | FBEa$^{*m}$ (n=52)<br>vs control (n=40) | 0-10<br>70-80<br>90-100 | <0.001<br>0.644<br>0.751 | ***<br>ns<br>ns |
| 3E[2] | FBEa$^{m}$ (n=64)<br>vs control (n=37) | 0-10<br>70-80<br>90-100 | <0.001<br>0.365<br>0.416 | ***<br>ns<br>ns |
| 3D, E | FBEa$^{m}$ (n=64)<br>vs FBEa$^{m}$ FBEa$^{*m}$ (n=26) | 0-10<br>70-80<br>90-100 | 0.663<br>0.001<br><0.001 | ns<br>**<br>*** |
| 3F | FBEa$^{m}$ FBEa$^{*m}$ (n=26)<br>vs control (n=20) | 0-10<br>70-80<br>90-100 | <0.001<br>0.036<br>0.006 | ***<br>*<br>** |
| 3G | FBEa$^{*m}$ FBEb$^{m}$ (n=49)<br>vs control (n=37) | 0-10<br>70-80<br>90-100 | 0.018<br>0.025<br>0.009 | *<br>*<br>** |
| 3D, G[3] | FBEa$^{*m}$ FBEb$^{m}$(n=49)<br>vs FBEa$^{*m}$ (n=31)[2] | 0-10<br>70-80<br>90-100 | 0.568<br>0.007<br>0.002 | ns<br>**<br>** |
| | **Reporter** | | | |
| S3B | FBEa$^{m}$ (n=24) vs wt (n=25)[2] | 0-10<br>70-80<br>90-100 | <0.001<br>0.693<br>0.350 | ***<br>ns<br>ns |
| S3B | FBEa$^{m}$ FBEa$^{*m}$ (n=19)<br>vs wt (n=25) | 0-10<br>70-80<br>90-100 | <0.001<br>0.010<br>0.003 | ***<br>*<br>** |

[1] Significance: *** < 0.001; ** < 0.01; * < 0.05; ns > 0.05

[2] Endogenous FBEa data from Carrick et al, 2024. Reporter FBEa data from this work.

[3] Separate data set from Figure 3D, done in same experiment as FBEa$^{*m}$ FBEb$^{m}$

**Supplementary Table 3**. Cryo-EM data collection and processing.

| | 1 FBF-2 | 2 FBF-2 |
|---|---|---|
| EMDB code | EMD-45096 | EMD-45097 |
| Magnification | 45,000 | |
| Voltage (kV) | 200 | |
| Electron exposure ($e^{-}$/Å$^{2}$) | 54 | |
| Defocus range (µm) | -1.0 to -2.5 | |
| Pixel size (Å) | 0.932 | |
| Symmetry imposed | C1 | |
| Initial particle images (no.) | 1,263,785 | |
| Final particle images (no.) | 252,126 | 110,843 |
| Map resolution (Å) (FSC = 0.143) | 4.4 | 6.4 |

**Supplementary Table 4.** Table of closely spaced adjacent FBF-2 binding sites identified in analysis of eCLIP data.

See enclosed Excel spreadsheet.

**Supplementary Table 5.** RNA sequences used in the EMSA RNA-binding experiments.

| **RNA** | **Sequence (with 3´-Cy5)**[1] |
|---|---|
| FBEa-FBEa* | AUCA**UGU**GCCAUACAUCA**UGUU**GCCAUUU |
| FBEa$^{m}$-FBEa* | AUCAACAGCCAUACAUCA**UGUU**GCCAUUU |
| FBEa-FBEa*$^{m}$ | AUCA**UGU**GCCAUACAUCAACA**U**GCCAUUU |
| FBEa$^{m}$-FBEa*$^{m}$ | AUCAACAGCCAUACAUCAACA**U**GCCAUUU |
| FBEa* | AUCA**UGUU**GCCAUUU |

[1]Mutations in red. FBE sequences underlined.

**Supplementary Table 6.** Sequences of guide RNAs and repair templates to create CRISPR alleles.

| **Description** | **Guide (5´-3´)** | **Repair template (5´-3´)** |
|---|---|---|
| FBEa | aaaaatggcaacatgatgta | gttcgttctcaccattttaggttaccatagaatcaACAgcGatacatcatgttgccattttttccccctctcatctcccc |
| FBEa* | aaaaatggcaacatgatgta | gttcgttctcaccattttaggttaccatagaatcatgtgccatacatcaACAtgccattttttccccctctcatctcccc |
| FBEa-FBEa* | aaaaatggcaacatgatgta | gttcgttctcaccattttaggttaccatagaatcaACAgcGatacatcaACAtgccattttttccccctctcatctcccc |
| FBEb | ataacTGTgaaaaataaagg | cccattcatactacctcgaatgccaaagcacccccctttattttttcACAgttatcttaacgctaacccctgtagaatcttcccggt |

**Supplementary Table 7.** Strains used in this manuscript.

| | **Strain name** | **allele** | **Comments** |
|---|---|---|---|
| N2 | | | |
| *sur-5* | JK4864 | *qIS147* | sur-5::GFP marked wt control |
| *unc-119(ed3) III; teIs1 IV* | TX189 | *teIs1* | oma-1::GFP prevents GFP silencing |
| **Endogenous *gld-1* Crispr alleles** | | | **Oligos to detect FBE mutations (5´-3´)** |
| *gld-1* FBEa$^{m}$ [1] | JK6540 | *q1242* | slc299 GAAGTACCCAACAACCACTTCG<br>prHJS401 TGGCAACATGATGTATCGCTGT |
| *gld-1* FBEa*$^{m}$ | JK6531 | *q1234* | slc299 GAAGTACCCAACAACCACTTCG<br>slc301 GAGAGGGGGAAAAAATGGCATGT |
| *gld-1* FBEa$^{m}$-FBEa*$^{m}$ | JK6541 | *q1243* | Use primer sets for FBEa and a* |
| *gld-1* FBEa*$^{m}$-FBEb$^{m}$ | JK6736 | *q1297* | Use primer set for FBEa*<br>For FBEb:<br>slc299 GAAGTACCCAACAACCACTTCG<br>slc302 GGGTTAGCGTTAAGATAACTGT |
| **Reporter strains and Crispr alleles** | | | |
| *rajSi50* FBEwt [2] | NIK50 | *rajSi50* | slc314 GTCACCAAGTACACTTCCAGCAAG<br>slc301 GAGAGGGGGAAAAAATGGCATGT |
| *rajSi50* FBEa$^{m}$ [1] | JK6551 | *q1274* | slc314 GTCACCAAGTACACTTCCAGCAAG<br>prHJS401 TGGCAACATGATGTATCGCTGT |
| *rajSi50* FBEa$^{m}$-FBEa*$^{m}$ | JK6639 | *q1275* | Use primer sets for FBEa and a* |

[1]Carrick et al. [2] Theil et al.

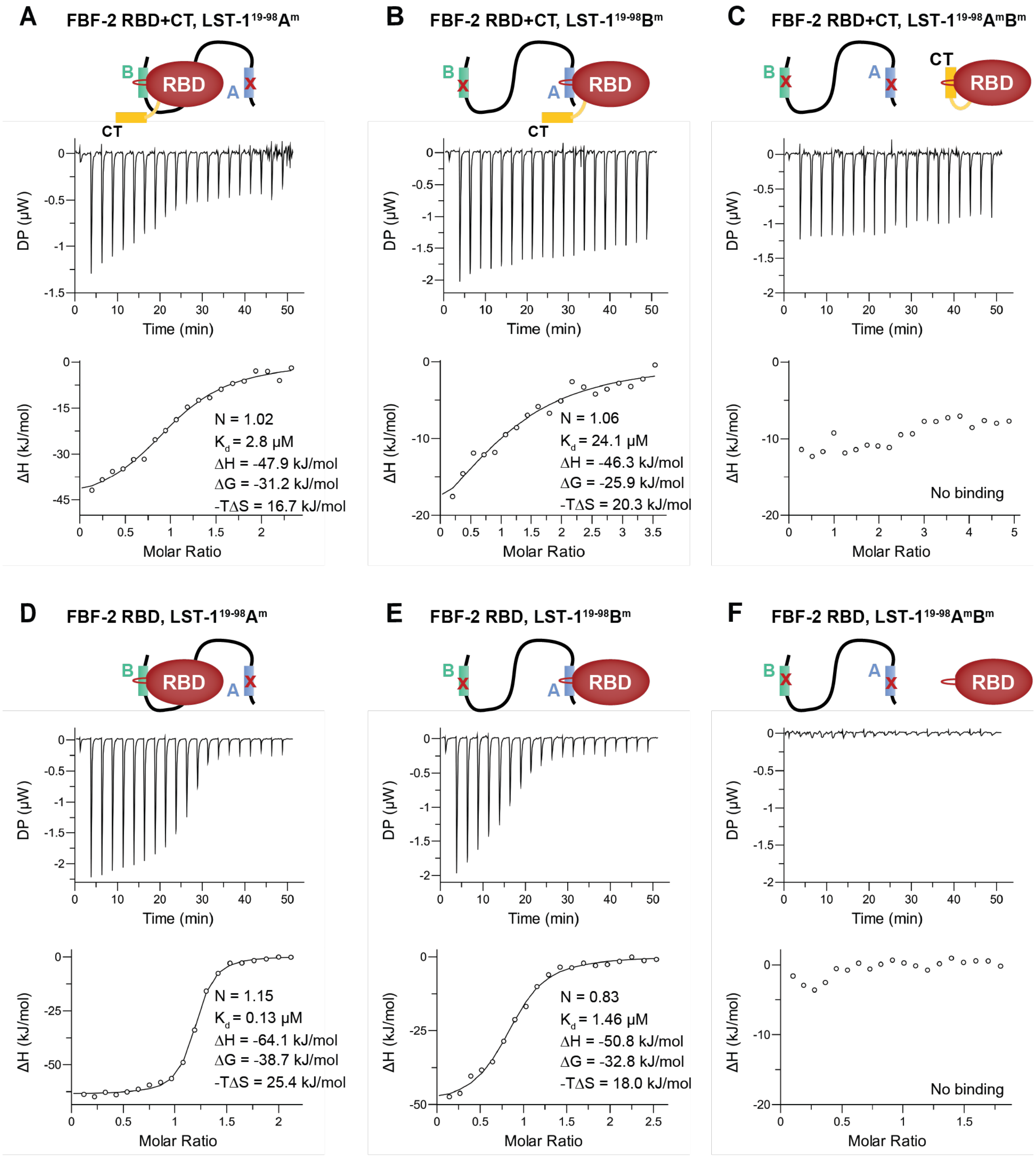


**Supplementary Figure 1.** LST-1 requires both PIMs for binding to two FBF-2 molecules. Representative ITC thermograms (top, differential power [DP] vs time) and corresponding titration curve-fitting graphs (bottom) for interaction of FBF-2 RBD+CT and (A) LST-$1^{19\text{-}98}$($A^m$), (B) LST-$1^{19\text{-}98}$($B^m$), and (C) LST-$1^{19\text{-}98}$ ($A^mB^m$). Representative ITC thermograms (top, DP vs time) and corresponding titration curve-fitting graphs (bottom) for interaction of FBF-2 RBD and (D) LST-$1^{19\text{-}98}$($A^m$), (E) LST-$1^{19\text{-}98}$($B^m$), and (F) LST-$1^{19\text{-}98}$($A^mB^m$). Thermodynamic parameters from one replicate indicated in bottom panels; thermodynamic parameters from two technical replicates are presented in **Table 1**. Experimental components indicated in diagrams above graphs.

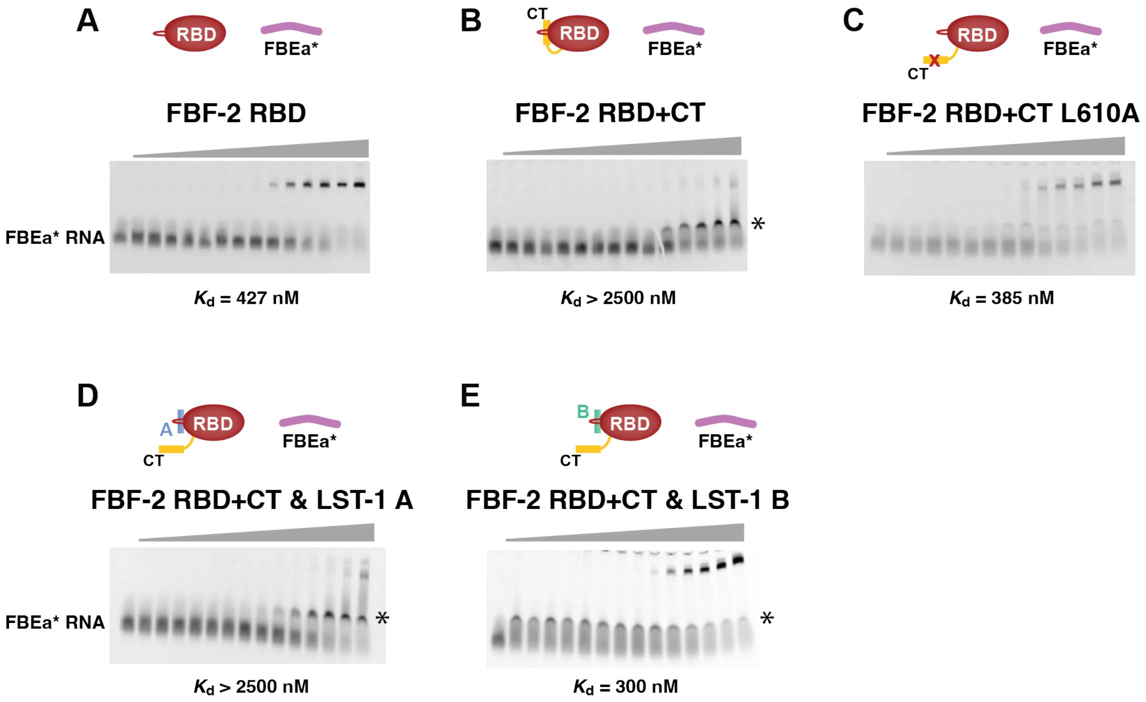


**Supplementary Figure 2.** FBF-2 binds to FBEa* RNA. Representative EMSA gels are shown for binding to FBEa* RNA (5´-AUCAUGUGCCAUAC-3´) by (A) FBF-2 RBD, (B) FBF-2 RBD+CT, (C) FBF-2 RBD+CT L610A, (D) FBF-2 RBD+CT with 150 µM LST-$1^{19\text{-}50}$ carrying PIM A, and (E) FBF-2 RBD+CT with 50 µM LST-$1^{67\text{-}98}$ carrying PIM B. Experimental components indicated in diagrams above gels. In panels B and D, we observed an intermediate band (*) that appears to be a non-specific interaction of FBF-2 RBD+CT, which was not observed for RBD. Similarly, LST-$1^{67\text{-}98}$ binds non-specifically to the RNA in panel E. We previously identified similar bands for non-specific binding to shorter RNAs[1]. We included these bands as part of the unbound RNA. Mean $K_d$ values from at least three technical replicates are reported. See also **Figure 2F**.

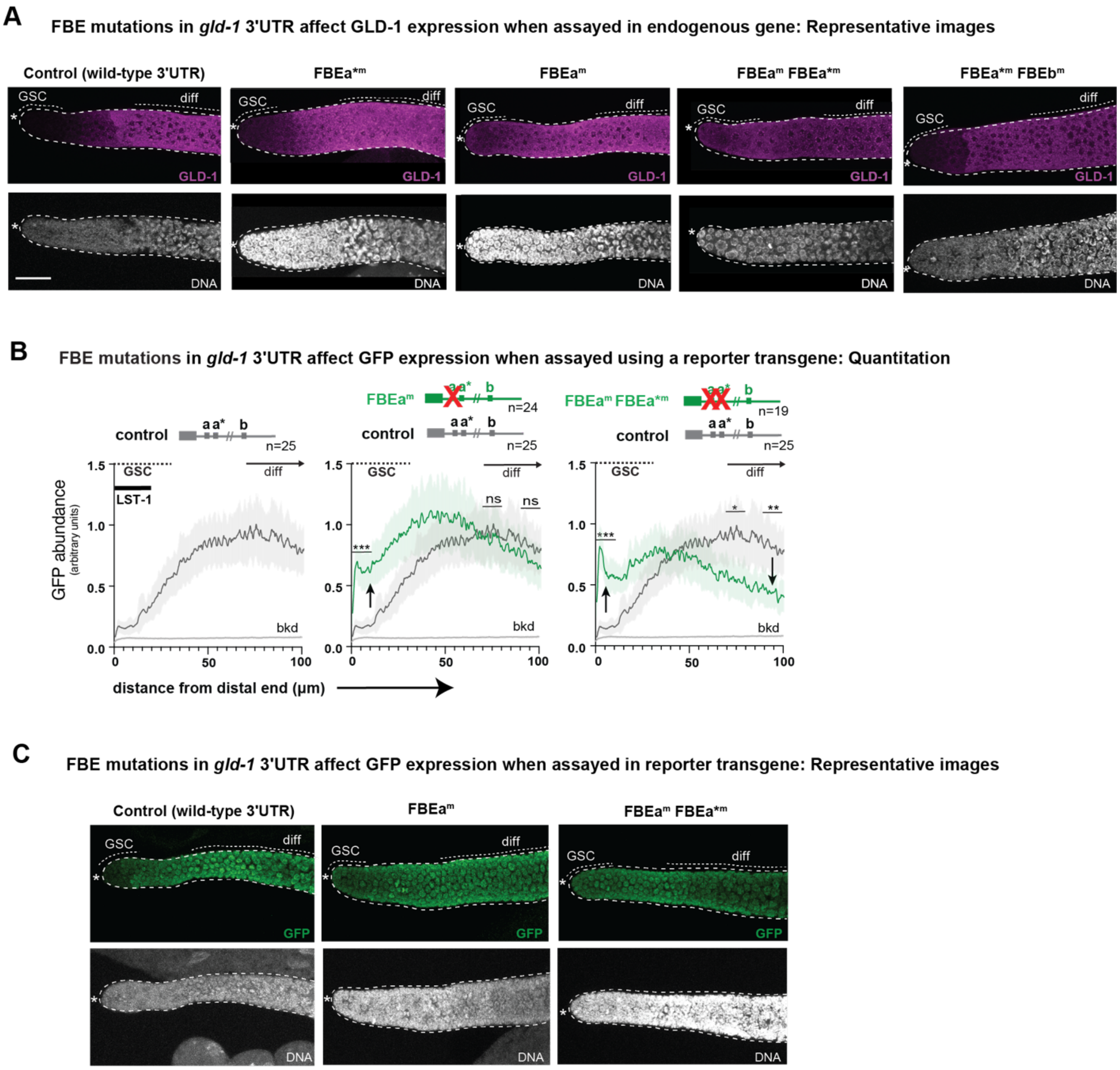


**Supplementary Figure 3.** Supplementary information for *gld-1* FBE mutants. (**A**) Representative z-projections of GLD-1 staining of FBE mutations generated in endogenous *gld-1*. Left to right: Thin dotted lines mark GSC pool and differentiation (diff); thick dotted line marks gonad boundary; asterisk marks distal end. Scale bar (bottom left), 20 µm for all panels in A and C. See Figure 3D-G for quantitation. See Carrick et al.[2] for images of GLD-1 in FBEa$^{m}$ mutant (graph in Figure 3E). (**B**) ImageJ quantitation of GFP abundance expressed from a *gld-1* 3´UTR reporter transgene as a function of position in the distal gonad. Gray lines show GFP pattern in wild-type; green lines show GFP pattern in mutant. Shading is the 95% confidence interval.

Gonadal regions with GSCs (GSC) and differentiated (diff) germ cells are marked above; extent of LST-1 protein is marked with a thick black line in the control panel. P-values are given for pooled data in 0-10, 70-80 and 90-100 µm regions (black bars). P-values: *** $p < 0.001$, ** $p < 0.01$, * $p < 0.05$, ns (not significant) $p > 0.05$. See **Supplementary Table 2** for exact p-values. Left to right: Wild-type control (two replicates), FBEa$^{m}$ (two replicates), and FBEa$^{m}$ FBEa*$^{m}$ double mutant (two replicates). Reporter constructs are shown above. Arrows indicate significant changes in GFP abundance relative to the wild-type control *gld-1* 3´UTR. Light grey line labeled “bkd” represents mean and 95% confidence interval for background staining in a wild-type animal not expressing the reporter construct (n=16, 1 replicate)[2]. (**C**) Representative z-projections of GFP reporter in extruded gonads with wild-type or mutated FBEs.

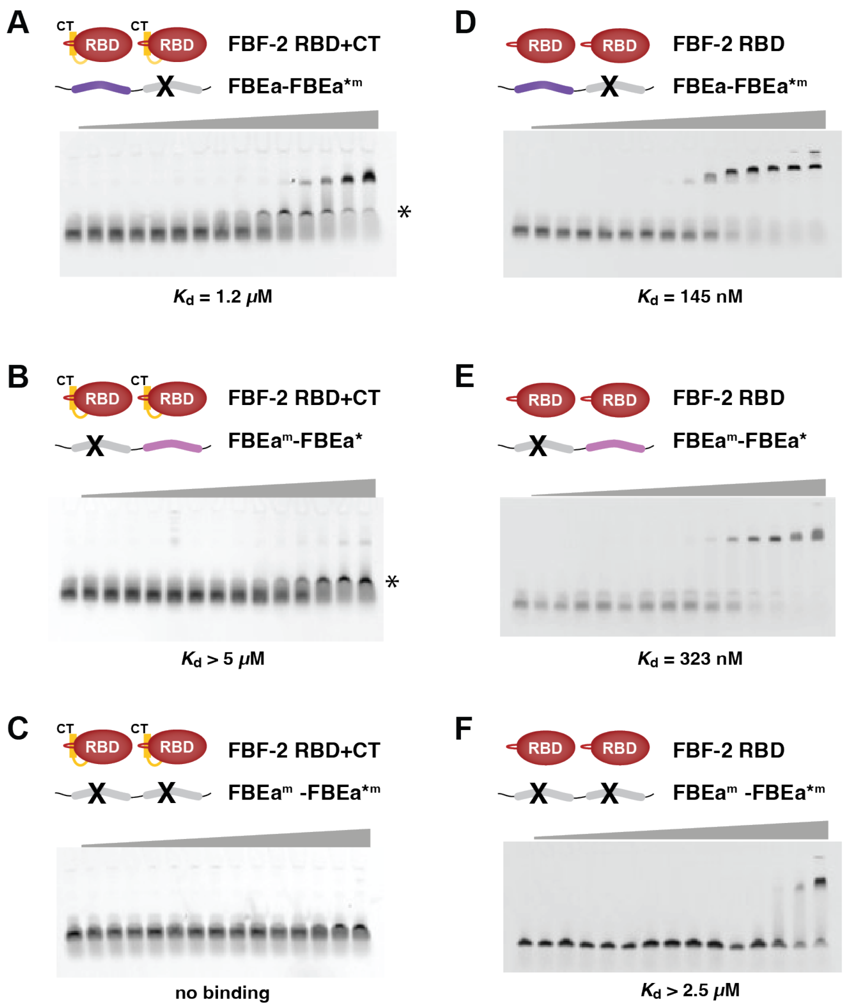


**Supplementary Figure 4.** Representative EMSA gels are shown for binding to FBEa-FBEa* RNA variants by (A-C) FBF-2 RBD+CT and (D-F) FBF-2 RBD. Experimental components indicated in diagrams above gels. LST-1 protein was not present. In panels A and B, we observed an intermediate band (*) that appears to be non-specific interaction of FBF-2 RBD+CT. We included these bands as part of the unbound RNA. Similar bands were detected previously with FBF-2 RBD+CT[1]. Mean $K_d$ values from at least three technical replicates are reported. See also **Table 2**.

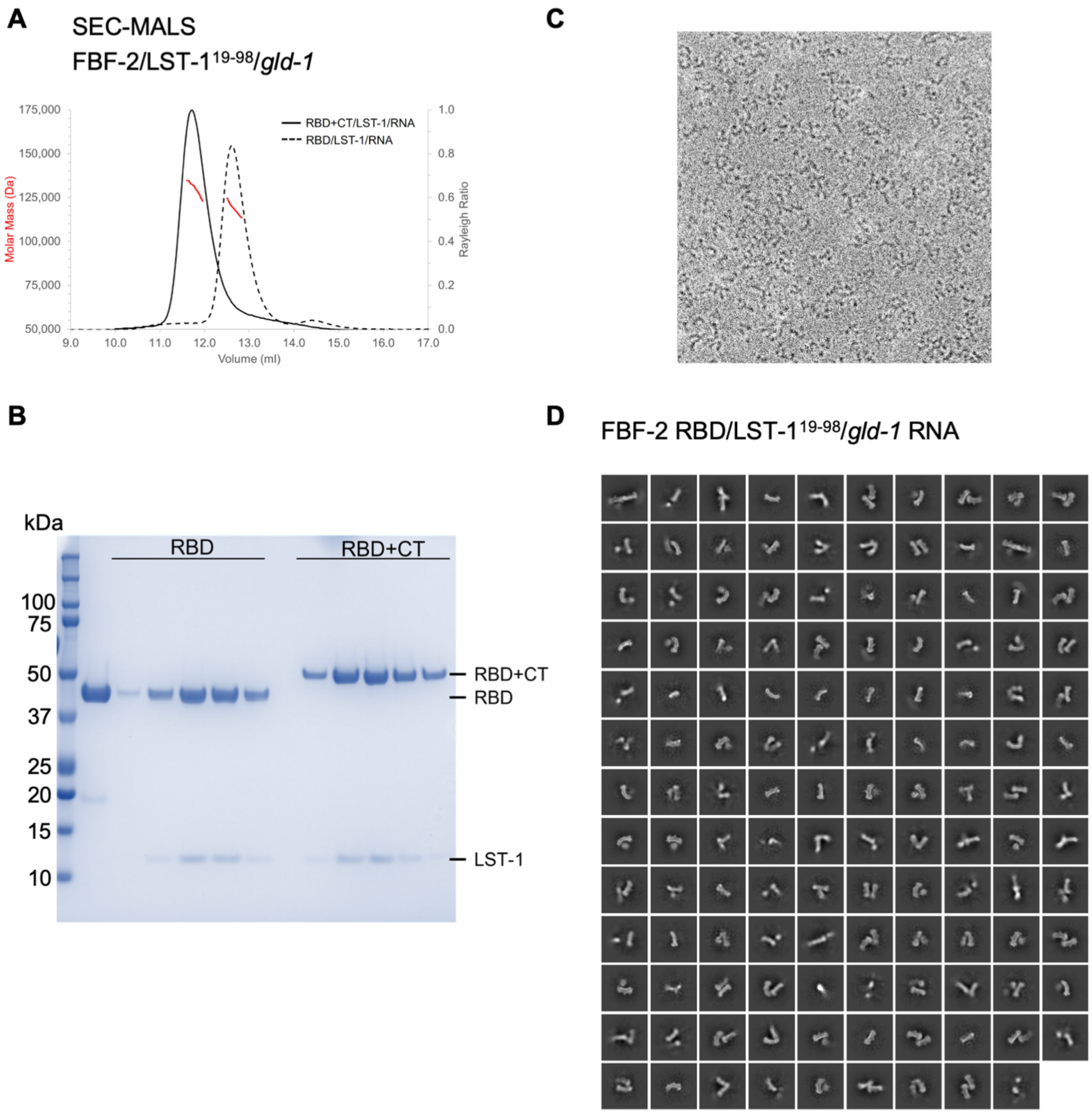


**Supplementary Figure 5.** Analyses of FBF-2/LST-1/ FBEa-FBEa* RNA complexes. (A) SEC-MALS analysis of FBF-2/LST-1$^{19\text{-}98}$/ FBEa-FBEa* RNA quaternary complexes. The peak for complexes with FBF-2 RBD+CT (solid black) had an apparent molecular mass of 130 kDa (red), which matches the calculated molecular weight of 125 kDa for a 2:1:1 complex of FBF-2 RBD+CT/LST-1$^{19\text{-}98}$/FBEa-FBEa*. The peak for complexes with FBF-2 RBD (dashed black line) had an apparent molecular mass of 119 kDa (red), which matches the calculated molecular weight of 119 kDa for a 2:1:1 complex of FBF-2/LST-1$^{19\text{-}98}$/FBEa-FBEa*. Molecular weights for individual components: FBF-2 RBD+CT, 53 kDa; FBF-2 RBD, 47 kDa; LST-1$^{19\text{-}98}$, 9.4 kDa, and

FBEa-FBEa* RNA, 8.6 kDa. (B) Coomassie-stained gel of SEC-MALS peak fractions. (C) Representative cryo-EM micrograph. (D) All 2D classes used for 3D reconstructions.

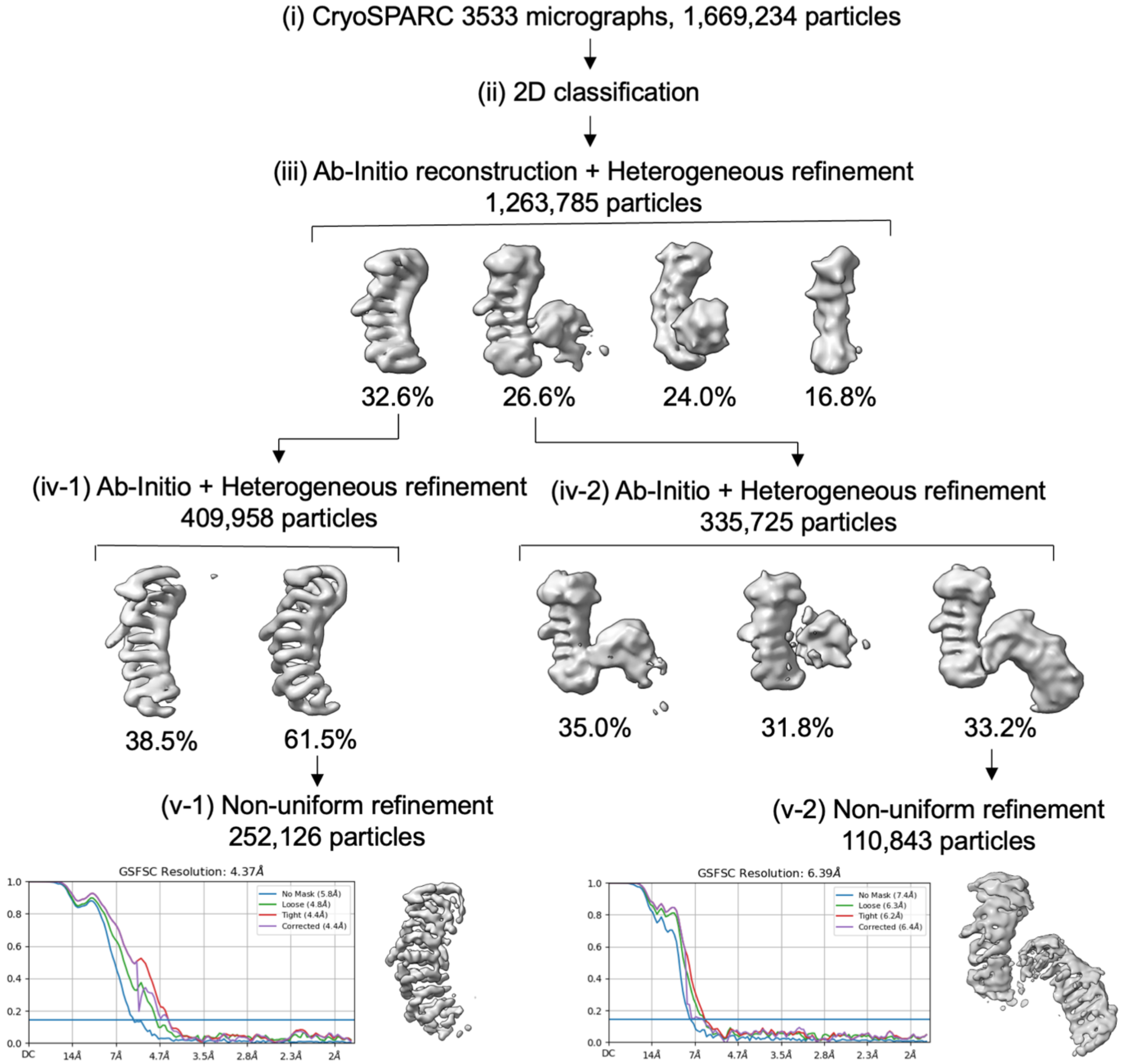


**Supplementary Figure 6.** Overview of cryo-EM processing scheme of FBF-2/LST-1/ FBEa-FBEa* RNA complex. Details are described in the Methods.

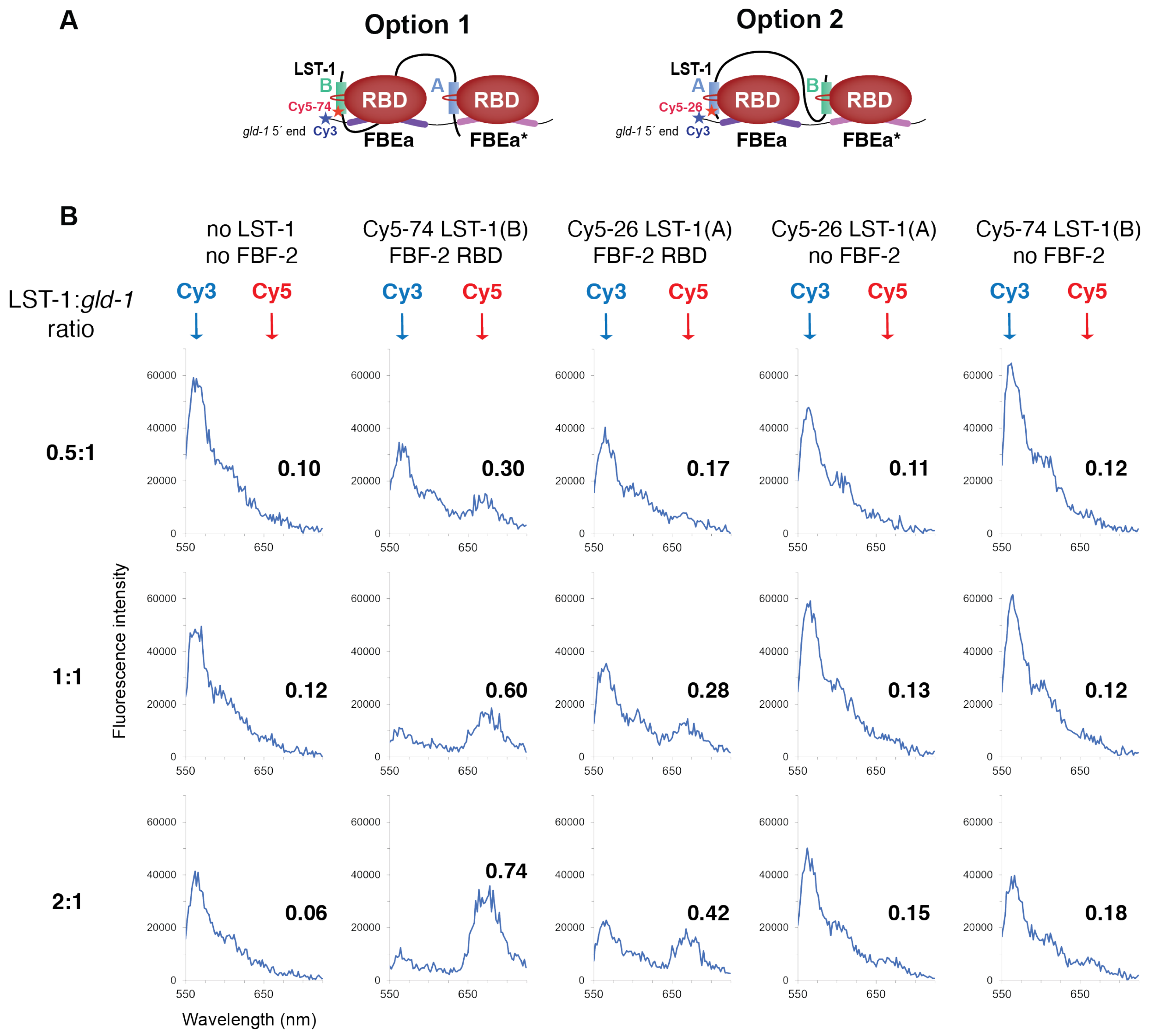


**Supplementary Figure 7.** FRET analysis of FBF-2/LST-1/ FBEa-FBEa* RNA complex. (A) Two possible orientations of LST-1 in the FBF-2/LST-1/ FBEa-FBEa* RNA complex. (B) Fluorescence spectra of excitation of Cy3-labeled RNA (blue arrow) and transfer to Cy5-labeled LST-1 (red arrow). In addition to complexes composed of Cy5-labeled LST-1, unlabeled FBF-2 RBD, and 5´-Cy3 labeled *gld-1* FBEa-FBEa* RNA, we also measured background levels of emission at 668 nm with samples of RNA only or RNA with LST-1 in the absence of FBF-2. FRET efficiencies were calculated as $I_{668}/(I_{668} + I_{564})$.

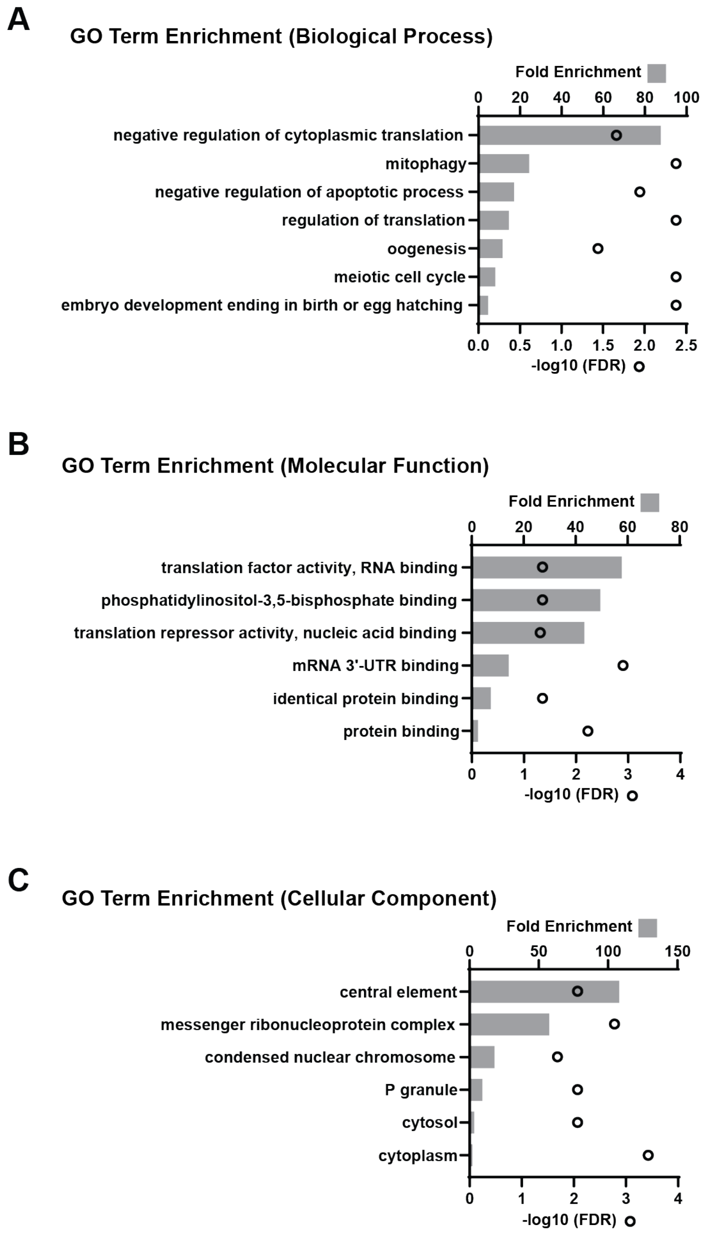


**Supplementary Figure 8.** GO term analysis of RNAs with adjacent FBEs. (A) Biological process GO term enrichment of genes that contain a peak with adjacent sites. Bars represent fold enrichment over background (top x-axis). Dots represent false discovery rate (FDR, bottom x-axis). Cutoffs for GO terms: fold enrichment ≥ 2 and FDR ≤ 0.05. (B) Molecular function GO term enrichment. Conventions and cutoffs as in (A). (C) Cellular component GO term enrichment. Conventions and cutoffs as in (A).
